# Absolute quantitative proteomics guides patient-stratified drug repurposing in clear cell and papillary renal cell carcinoma

**DOI:** 10.64898/2026.07.17.738718

**Authors:** André Q. Figueiredo, Inês F. Domingos, Carlos Lodeiro, Rajiv Dhir, Luís B. Carvalho, Laura Mercolini, Jacek R. Wiśniewski, Gianluca Moroncini, Mariana Medeiros, Luís Campos Pinheiro, Hélder Mansinho, Hugo M. Santos, Dimitrios Korentzelos, José L. Capelo

**Author notes:** Authors to whom correspondence should be addressed: José L. Capelo, Dimitrios Korentzelos.

## Abstract

**Background:** Renal cell carcinoma (RCC) is a highly heterogeneous disease in which distinct molecular subtypes exhibit characteristic genomic, metabolic, and microenvironmental features that influence therapeutic response. Substantial inter-patient variability exists within each subtype, resulting in markedly different clinical outcomes even among tumours of the same histological category. Proteomics provides a direct readout of tumour biology and pathway activity, complementing genomic information and enabling the identification of patient-specific actionable vulnerabilities. We applied a Total Protein Approach (TPA)-based prescriptomics framework that integrates absolute quantitative proteomics with curated drug–target knowledge to nominate patient-specific drug-repurposing options, positioned as coadjuvants to the prevailing standard of care.

**Methods:** Seventeen human kidney tissue specimens, seven clear cell RCC (ccRCC), five papillary RCC (pRCC), and five normal adjacent tissues (NAT), were retrieved from the publicly available PRIDE repository (PXD023296) and reanalysed by TPA-based absolute quantification applied to previously acquired label-free LC-MS/MS data. Differential expression analysis between each tumour subtype and NAT identified subtype-specific upregulated proteins, wich were intersected with Therapeutic Target Database (TTD) to nominate FDA-approved drugs targeting dysregulated proteins as candidate repurposing strategies.

**Results:** ccRCC and pRCC produced distinct proteome-wide upregulation profiles consistent with their known biological drivers. TPA index stratification nominated bempedoic acid (ACLY inhibitor) and tipiracil hydrochloride (TYMP inhibitor) as patient-stratified candidates for ccRCC, and auranofin (TXNRD1 inhibitor), bempedoic acid, and mipomersen (APOB-directed antisense oligonucleotide) for pRCC. ACLY was the only top-priority target shared across both subtypes, pointing to a candidate cross-subtype metabolic vulnerability. Secondary candidates emerged from protein–protein interaction network analysis in both subtypes..

**Conclusions:** This study presents a quantitative proteomics framework for translating individual-patient proteomic dysregulation into coadjuvant drug-repurposing hypotheses across the principal RCC subtypes. By combining the TPA for absolute protein quantification with prescriptomics-guided drug–target mapping, we show that ccRCC and pRCC harbour distinct, individually stratifiable therapeutic vulnerabilities. These findings provide a proof-of-concept for proteomics-based treatment stratification in RCC and establish a scalable framework that, pending functional validation, could inform personalised therapeutic decision-making across RCC subtypes.

## Introduction

Renal cell carcinoma (RCC) represents approximately 3% of adult malignancies and is the most common type of kidney cancer, arising from the renal tubular epithelium^1,2^. RCC comprises a heterogeneous group of tumours with distinct histopathological features, genetic alterations, biological behaviour and therapeutic responsiveness^3^. Its principal subtypes, clear cell RCC (ccRCC) and papillary RCC (pRCC) differ in molecular drivers, clinical course and treatment sensitivity, yet both are managed by histological category rather than individual tumour biology^3,4^.

Clear cell RCC is the most prevalent subtype, accounting for 70–75% of RCC cases, and is defined histologically by cells with clear cytoplasm due to glycogen and lipid accumulation^5^. The molecular hallmark of ccRCC is inactivation of the Von Hippel–Lindau (VHL) tumour suppressor gene, observed in up to 90% of cases, leading to stabilisation of hypoxia-inducible factors (HIF-1α and HIF-2α) and upregulation of angiogenic and metabolic programmes, including VEGF, PDGF and CAIX^6^. Treatment of advanced ccRCC has evolved considerably over the past two decades, from tyrosine kinase inhibitors (TKIs) targeting VEGFR, exemplified by sunitinib^7^, to single-agent immune checkpoint inhibitors (ICIs) such as nivolumab^8^, and more recently to dual PD-1/CTLA-4 blockade with nivolumab plus ipilimumab^9^ and to first-line Immune Checkpoint Inhibitors (ICI)–TKI combinations including pembrolizumab plus axitinib^10^ and nivolumab plus cabozantinib^11^.

Adjuvant TKI therapy has also been explored in resected high-risk disease^12^. Although these regimens extend progression-free and overall survival, response remains heterogeneous, and most patients ultimately progress, underscoring the need for molecularly guided personalisation^13^.

Papillary RCC accounts for approximately 10–20% of RCCs. Historically subdivided into type 1 and type 2, papillary RCC is now recognised as a heterogeneous entity, and the 2022 WHO classification no longer recommends this simple binary subtyping. Instead, several tumours previously grouped within type 2 papillary RCC are now classified as distinct entities, including FH-deficient RCC, a molecularly defined and aggressive renal carcinoma frequently associated with hereditary leiomyomatosis and RCC (HLRCC)^4,14^. Tumours historically classified as type 1 papillary RCC are generally less aggressive and are enriched for Mesenchymal-epithelial transition (MET) pathway activation through mutation, amplification or chromosomal gains^14^. MET inhibitors such as savolitinib have demonstrated clinical activity in MET-driven papillary RCC^15^. Fumarase hydratase (FH)-deficient RCC is characterised by fumarate accumulation, pseudohypoxic signalling, metabolic rewiring and epigenetic dysregulation^16–18^.

Proteomics-based analysis enables functional characterisation of RCC beyond genomics: unlike DNA or RNA profiling, proteomics captures dynamic features such as protein expression levels, post-translational modifications and pathway activity^19^. Several studies have combined high-resolution mass spectrometry with the Total Protein Approach (TPA) to identify novel biomarkers and druggable targets in RCC^20,21^. The TPA enables label- and standard-free estimation of absolute protein abundances by normalising each protein’s signal to the total proteome signal, making it cost-effective and scalable for large or exploratory studies^22,23^. Its accuracy, however, depends on comprehensive proteome coverage, and missing or poorly detected proteins can bias normalisation^22,23^. Inter-laboratory variability and differences in digestion, ionisation and instrument response may impair comparability, often requiring complementary strategies such as reference protein normalisation^23,24^For applications demanding precise absolute quantification, isotope-labelled standards or validated immunoassays remain preferable^24^. Nonetheless, TPA performs well for biomarker discovery and longitudinal analyses when carefully applied^20,21,25^. For instance, TPA-based profiling identified subtype-specific protein signatures, PLIN2 for ccRCC and TUBB3 for pRCC, that were subsequently validated by immunohistochemistry^21^.

Prescriptomics, the integration of omics-derived molecular profiles with curated drug–target knowledge to nominate patient-specific therapies, has emerged as a promising precision oncology concept^26^. Drug repurposing offers a pragmatic route to expand cancer therapeutics: repositioning compounds with established safety profiles and regulatory history can shorten development timelines and reduce costs relative to *de novo* drug discovery^26^. By integrating TPA-based proteomics with curated drug–target databases, differentially expressed proteins can be mapped to clinically approved drugs or investigational compounds, enabling patient-level therapeutic prioritisation^27^. For example, in ccRCC, activation of hypoxia, angiogenic, glycolytic and PI3K/AKT/mTOR-related programmes provides a molecular rationale for targeting VEGFR- and mTOR-associated pathways^28^. In MET-driven pRCC, integrative molecular profiling of the TCGA papillary cohort supports the mechanistic basis for MET-targeted therapy^29^. In FH-deficient tumours, fumarate accumulation, redox imbalance and metabolic rewiring suggest potential vulnerabilities involving mitochondrial metabolism, oxidative stress and related adaptive pathways^30,31^. Resources such as the Therapeutic Target Database (TTD)^27^ and large-scale pharmacogenomic resources such as the Genomics of Drug Sensitivity in Cancer database^32^ complement this strategy by linking molecular alterations to candidate therapeutic agents. This is particularly relevant in RCC, where pronounced intratumour heterogeneity and resistance to single-agent therapies limit the durability of monotherapy^33^.

In this work, we applied a TPA-based prescriptomics framework to 17 human kidney tissue specimens spanning the main principal RCC subtypes, ccRCC and pRCC, together with normal adjacent tissue (NAT). Our work presents two main contributions. First, we use absolute, rather than relative, protein quantification to rank candidate drug targets, enabling within-patient prioritisation via a TPA-based index (TPA index). Second, we perform a personalisation at the patient level, rather than the subtype level, so that two patients with identical histological diagnoses may receive distinct nominated therapies. Together, these features establish a proof of concept for proteomics-guided basis for individualised drug-repurposing hypotheses in RCC and a template that could be extended to other heterogeneous malignancies.

## Methods

### Study design and sampling

In the present work, we re-analysed a previously published proteomics dataset comprising 17 flash-frozen, optimal cutting temperature (OCT)-embedded human renal tissue specimens^20,34,35^ (PRIDE: PXD023296). TPA-based absolute protein quantification was applied to derive patient-level TPA index values and to map upregulated proteins onto a curated drug–target database. These comprise clear cell renal cell carcinoma (ccRCC, *n* = 7), papillary renal cell carcinoma (pRCC, *n* = 5), and treatment-naïve normal adjacent tissue (NAT, *n* = 5) ^34^. All samples were sourced from the University of Pittsburgh Biospecimen Core, with ethical clearance granted by the University of Pittsburgh Institutional Review Board (IRB #02-077). Each tumour specimen contained a minimum of 85% neoplastic cells. Clinical details for the analysed patients are provided in Supplementary Table 1.

### Data analysis, protein-protein interaction, and dysregulated pathway analysis

The data processing pipeline for this work is presented in Fig. 1b. Data analysis and statistics were performed using a label-free quantification workflow implemented in MaxQuant V1.6.0.16, with all raw files processed under default settings^37,38^. Protein identification was achieved through the Andromeda search engine using the UniProt-SwissProt Human database and a common contaminants database. Prior to data processing, reverse hits and proteins identified only by site were removed, and TPA-based concentrations for each protein in the dataset were calculated using the formula shown in Fig. 1a (Equations 1 and 2) and detailed in Supplementary Material 1. The principle of TPA relies on the assumption that the total spectral intensity from all detected peptides in a mass spectrometry (MS)-based experiment is directly proportional to the concentration of the protein in the solution, as per equation (1)^22^ (Fig.1a, equation (1))

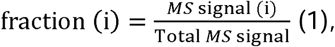

**Fig. 1.**
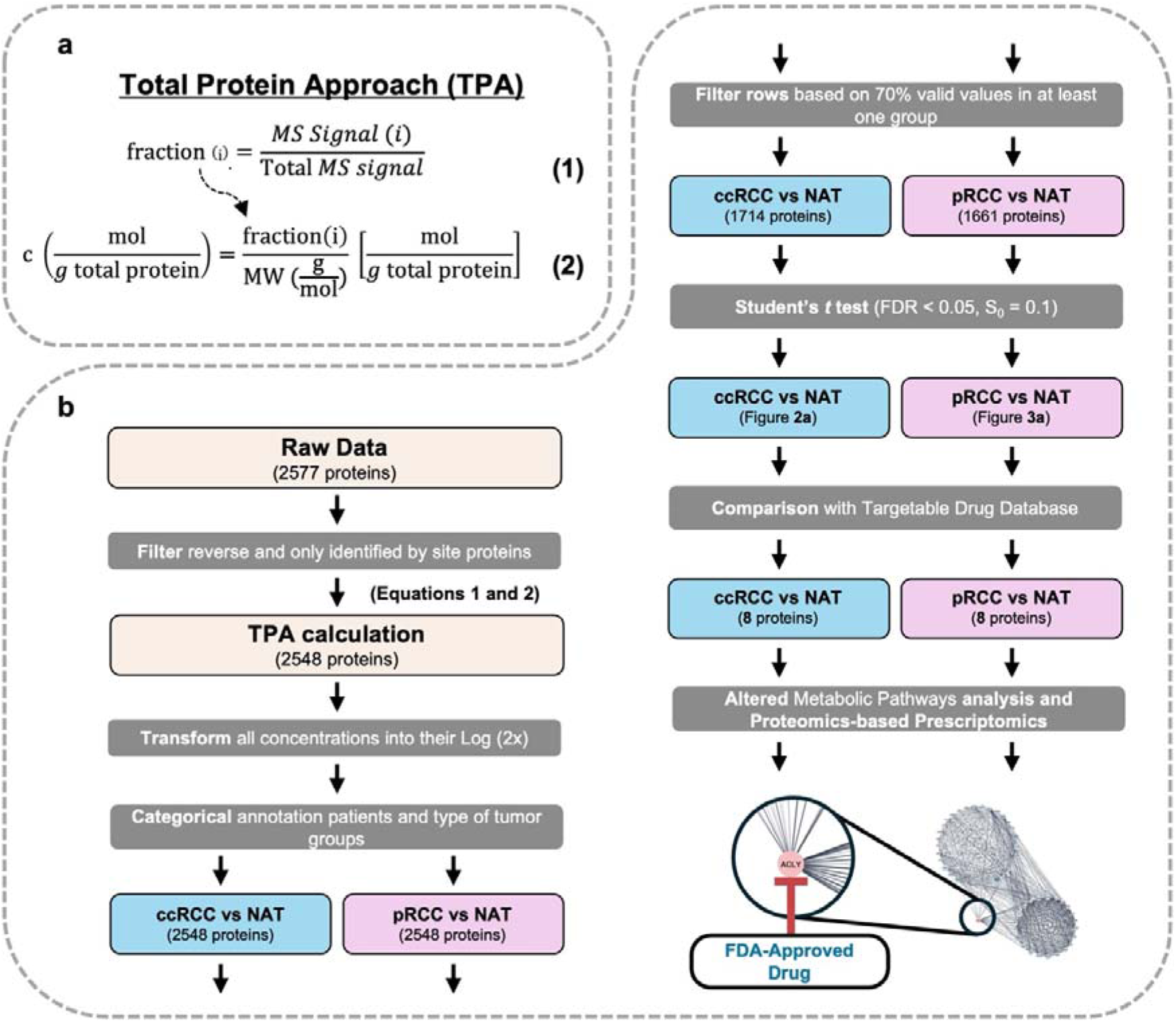
TPA-Based Quantitative Proteomics Workflow for Prescriptomic Target and Pathway Analysis in Renal Tumours. (**a**) Equations applied in the Total Protein Approach (TPA) to determine the concentration of each protein in the dataset (Equations 1 and 2). (**b**) Schematic representation of the proteomics workflow, including data processing, differential expression analysis, drug–target matching, and pathway analysis, used to derive patient-specific prescriptomic strategies based on proteomics for each renal neoplasm subtype (ccRCC and pRCC). To identify the most altered proteins in each cancer subtype, the TPA-based concentration of each protein was compared with the mean concentration in NAT, yielding the TPA index value (Equation 3). This TPA-based drug repurposing framework supports the targeting of individual proteins (ccRCC or pRCC).

where *MS*signal (i) corresponds to the spectral intensity of protein, and the Total *MS* signal is the sum of all protein spectral intensities in each sample ^22^. The protein molecular weight is then used to convert this relative signal into an absolute concentration in molar units (mol of protein per g of total protein), according to equation (2) (Fig. 1a, equation (2))^22^.

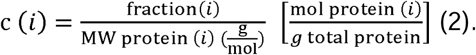

Consequently, for each protein with a measurable spectral intensity, a TPA-based concentration was calculated. Processed TPA-based concentration data were further analysed using Perseus V2.0.11 with default parameters^36,37^. TPA-based concentrations were further log_2_-transformed to minimise the influence of outliers and reduce the data skewness. Protein groups were only retained if present in at least 70% of samples within one group between the renal neoplasm-NAT comparison. The list of differentially expressed proteins was obtained by comparing two groups, a renal neoplasm subtype (ccRCC and pRCC) against NAT, using a two-tailed Student’s t-test (permutation-based FDR 0.05 and S0 of 0.1), and the log_2_ ratios were calculated as the difference in average log_2_ TPA-concentration values between the tested conditions (Supplementary Material 1). For each comparison, the upregulated proteins were matched with a database of FDA (Food and Drug Administration)-approved targets^27^ to identify possible new therapies for those renal neoplasms. To establish a patient-level protein importance ranking, a TPA-based index was calculated for each protein in each cancer sample by comparing its TPA-based concentration in the tumour of a given patient with the average concentration of the same protein across all NAT samples, according to equation (3):

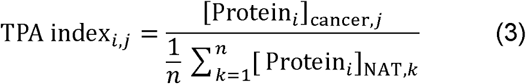

Where *i* denotes the protein, *j* denotes the cancer sample from an individual patient, and *n* is the number of NAT samples. Technical replicates were first averaged for each sample. Therefore, the numerator represents the mean TPA-based concentration of protein *i* in the cancer tissue of patient *j*, while the denominator represents the mean concentration of the same protein across all NAT samples. Proteins with a TPA index_i_,_j_>2 were considered overrepresented in the cancer sample of that patient and selected as candidates for further analysis. The TPA index threshold of 2 was selected *a priori* as an operational cutoff representing a minimum twofold overabundance in tumour relative to mean NAT concentration. This threshold was defined before data inspection. To assess the sensitivity of patient-level drug nominations to this choice, TPA index thresholds of 1.5, 3, and 4 were additionally evaluated; results are provided in Supplementary Figures 1 and 2. For each patient, the highest-ranked FDA-approved therapeutic target, as determined by the TPA index, was selected as the seed node for patient-specific protein-protein interaction (PPI) network analysis in Cytoscape v3.10.4 using the STRING v2.2.0 plugin. PPI networks for all cancer types were enriched using KEGG, Reactome, and WikiPathways as ontology databases to identify pathways enriched for FDA-approved drugs with established repurposing potential as coadjuvant therapies (Supplementary Material 1).

**Fig. 2.**
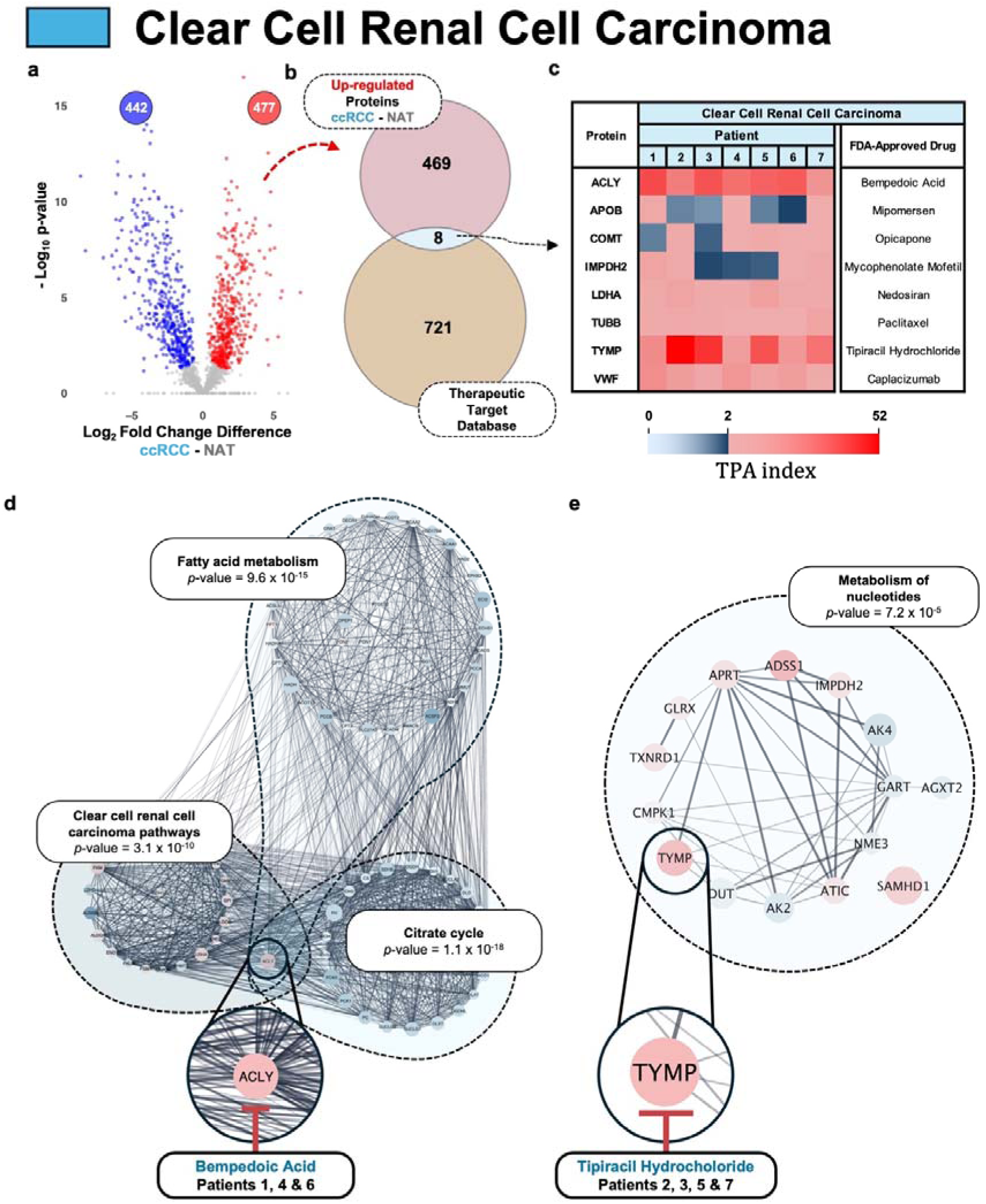
Patient-stratified prescriptomics in clear cell renal cell carcinoma (ccRCC). **(a)** Volcano plot of differential protein abundance in ccRCC versus normal-adjacent tissue (NAT) controls (log₂ fold change vs −log_10_ *p*-value). Proteins meeting the significance criteria (two-sided Student’s *t*-test, *p*-value < 0.05; *s*₀ = 0.1) are highlighted as upregulated (red; *n* = 477) or downregulated (blue; *n* = 442); non-significant proteins are shown in grey. (**b),** Intersection of upregulated proteins with the Therapeutic Target Database (TTD) identifies eight proteins with FDA-approved targeting opportunities (*n* = 8). (**c)** Heatmap of patient-level TPA index values (Equation 3) for the eight candidate drug targets across ccRCC patients (1–7); the colour scale denotes TPA index (0–2, blue gradient; >2–52, red gradient), with corresponding FDA-approved drugs indicated. (**d,e)** Functional network and pathway enrichment analysis (STRING/Cytoscape) for the phenotype-defining targets ACLY (**d**) and TYMP (**e**). ACLY-associated networks cluster in lipid and central carbon metabolism (fatty acid metabolism, *p*-value = 9.6 × 10^-15^; citrate cycle, *p*-value = 1.1 × 10^-18^), whereas TYMP-associated networks cluster in metabolism of nucleotides (*p*-value = 7.2 × 10^-15^). Putative repurposing hypotheses are prioritised by TPA index (bempedoic acid for ACLY-high: patients 1, 4 and 6; tipiracil hydrochloride for TYMP-high: patients 2, 3, 5 and 7) and are intended for experimental validation rather than clinical recommendation. In Cytoscape networks, nodes are coloured according to the volcano plot (red, upregulated; blue, downregulated) and scaled by significance (larger nodes indicate higher significance).

## Results

### Differential protein expression in ccRCC

Widespread proteomic remodelling in ccRCC relative to NAT was observed by volcano-plot analysis (Fig. 2a), with 477 proteins significantly upregulated and 442 downregulated in tumours. Intersecting the upregulated proteome with Therapeutic Target Database entries, followed by curation for FDA-approved drugs, identified eight targetable proteins (Fig. 2b). Patient-level TPA index values for these proteins, calculated using Equation 3, are summarised in Fig. 2c. Based on the highest-priority targetable proteins, the seven ccRCC patients were stratified into two molecular subgroups: three ACLY-high patients, corresponding to patients 1, 4 and 6, and four TYMP-high patients, corresponding to patients 2, 3, 5 and 7. Among the remaining targetable protein candidates, APOB, COMT, and IMPDH2 did not consistently exceed the TPA index threshold across patients, whereas LDHA, TUBB, and VWF (von Willebrand factor) showed elevated TPA index values in subsets of patients, supporting their prioritisation as secondary or tertiary therapeutic options. Network-based contextualisation of the top-ranked protein per patient using STRING/Cytoscape (Fig. 2d,e) demonstrated coherent pathway organisation. ACLY-linked networks were enriched for fatty-acid metabolism, citrate/TCA-cycle-related processes and ccRCC-associated pathways, whereas the TYMP-linked network was enriched for nucleotide metabolism. These protein- and pathway-level signals were integrated into the proposed sequential treatment framework for ccRCC (Fig. 4), in which first- to third-line candidates are prioritised according to their patient-specific TPA index values.

### Differential protein expression in pRCC

Volcano plot analysis identified 470 upregulated and 347 downregulated proteins in pRCC relative to NAT (Fig. 3a). Intersecting the upregulated proteome with the Therapeutic Target Database identified eight targetable proteins associated with FDA-approved drugs (Fig. 3b), for which patient-level TPA index values, calculated according to Equation 3, are shown in Fig. 3c. Among these candidates, ACLY showed the most consistent increase in absolute TPA-derived abundance, particularly in patients 2, 4 and 5, thereby defining the largest target-based subgroup. TXNRD1 and APOB also displayed markedly increased absolute abundance, but each predominated in a single patient, corresponding to patient 1 and patient 3, respectively. Thus, the five pRCC cases were stratified into three principal molecular profiles: TXNRD1-high, ACLY-high, and APOB-high. Among the remaining targetable proteins, LDHA and PARP1 showed the fewest TPA index values above the selection threshold, whereas PGD, TUBB and TYMP displayed increased absolute abundance in subsets of patients, supporting their prioritisation as secondary or tertiary therapeutic options. Network-based contextualisation of the top-ranked targetable proteins using STRING and Cytoscape revealed coherent pathway structures (Fig. 3d–f). TXNRD1-associated networks were enriched for cellular response to stress, nuclear events mediated by NFE2L2 and ferroptosis (Fig. 3d). ACLY-associated networks were primarily linked to altered metabolic processes, including fatty-acid metabolism and the citrate cycle (Fig. 3e). In the APOB-high case, enriched clusters involved plasma lipoprotein assembly, remodelling, and clearance, cholesterol metabolism, and fatty acids and lipoproteins transport (Fig. 3f). As observed for ccRCC, these protein- and pathway-level findings were integrated into the proposed sequential treatment framework for pRCC (Fig. 4), in which first-to third-line therapeutic options are prioritised according to patient-specific TPA index values.

**Fig. 3.**
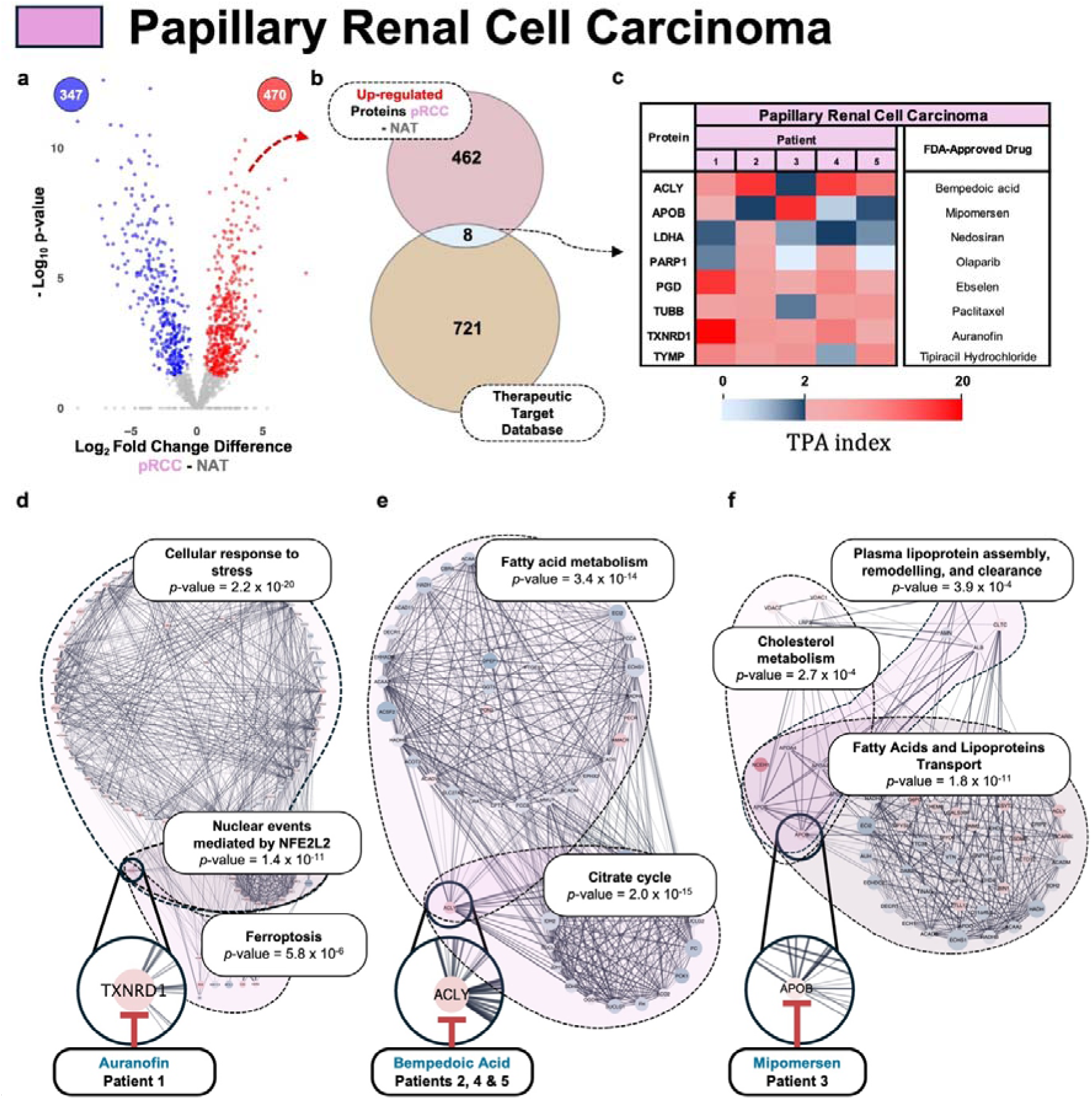
Patient-stratified prescriptomics in papillary renal cell carcinoma (pRCC). (**a**) Volcano plot of differential protein abundance in pRCC versus normal-adjacent tissue (NAT) controls (log₂ fold change vs −log10 *p*-value). Proteins meeting the significance criteria (two-sided Student’s *t*-test, *p*-value < 0.05; *s*₀ = 0.1) are highlighted as upregulated (red; *n* = 470) or downregulated (blue; *n* = 347); non-significant proteins are shown in grey. (**b**) Intersection of the upregulated pRCC proteome with the Therapeutic Target Database (TTD) identifies eight proteins with existing approved drugs annotated in TTD (*n* = 8). (**c**) Heatmap of patient-level TPA index values (Equation 3) for the eight candidate drug targets across pRCC patients (1–5); the colour scale denotes TPA index (0–2, blue gradient; >2–20, red gradient). (**d-f**) Functional network and pathway enrichment analysis (STRING/Cytoscape) for the phenotype-defining targets TXNRD1 (**d**), ACLY (**e**), and APOB (**f**). (**d**) TXNRD1-associated networks cluster in cellular response to stress, *p*-value = 2.2 × 10^-20^; nuclear events mediated by NFE2L2, *p*-value = 1.4 × 10^-11^; ferroptosis, *p*-value = 5.8 × 10^-6^). (**e**) ACLY-associated networks cluster in fatty acid metabolism, *p*-value = 3.4 × 10^-11^; citrate cycle, *p*-value = 2.0 × 10^-15^). (**f**) APOB-associated networks in plasma lipoprotein assembly, remodelling and clearance, *p*-value = 3.9 × 10^-4^; cholesterol metabolism, *p*-value = 2.7 × 10^-4^; fatty acid and lipoprotein transport, *p*-value = 1.8 × 10^-11^). Putative repurposing hypotheses are prioritised by TPA index (auranofin for TXNRD1-high: patient 1; bempedoic acid for ACLY-high: patients 2, 4, and 5; mipomersen for APOB-high: patient 3) and are intended for experimental validation rather than clinical recommendation. In Cytoscape networks, nodes are coloured according to the volcano plot (red, upregulated; blue, downregulated) and scaled by significance (larger nodes indicate higher significance).

**Fig. 4.**
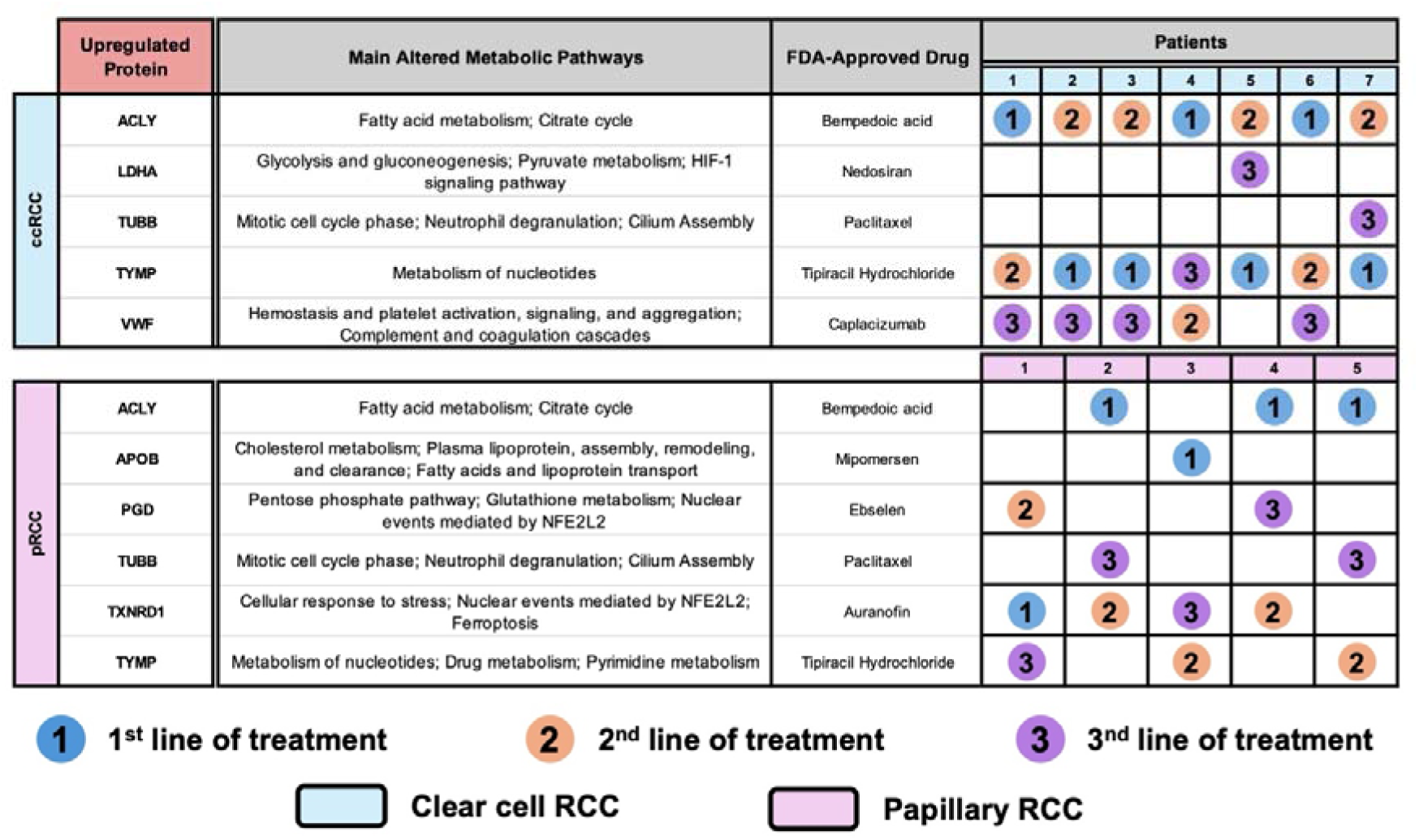
TPA-based prescriptomics nominates patient-stratified therapeutic hypotheses across RCC subtypes. Summary of the main upregulated targets (ccRCC and pRCC) identified by our TPA-based prescriptomics workflow, with associated dysregulated metabolic programmes and nominated repurposing candidates. The top-ranked upregulated proteins are shown (selected for readability from Figs. 2c and 3c), with priority rank based on TPA index stratification (1, highest; 2, second; 3, third). This figure summarises hypothesis-generating repurposing opportunities and does not imply established clinical lines of therapy. All candidates and lines of treatment require further experimental validation.

## Discussion

Applied across the two principal RCC subtypes, the TPA-based prescriptomics framework generated, patient-stratified therapeutic candidates from a single label-free LC–MS/MS experiment, without isotope labels, spike-in standards, or tumour-type-specific prior knowledge. In ccRCC and pRCC, proteome-wide upregulation yielded sufficient druggable signal for target-centric prioritisation via TTD intersection.

In ccRCC, our TPA index stratification highlighted two dominant, patient-separable signatures centred on ACLY and TYMP (Fig. 4). ACLY-high tumours showed enrichment of fatty acid metabolism and the citrate cycle, consistent with the oncogenic role of ACLY in cancer metabolism^38,39^. In this context, bempedoic acid, a first-in-class, oral ACLY inhibitor approved for hypercholesterolaemia and atherosclerotic cardiovascular disease^40,41^, emerged as a plausible candidate for experimental evaluation, given its potential to reduce cytosolic acetyl-CoA availability and thereby constrain lipogenic output. Because bempedoic acid is a prodrug requiring activation by very-long-chain acyl-CoA synthetase 1 (SLC27A2/ACSVL1), whose expression is highest in hepatocytes, tumour-specific target engagement cannot be assumed and should be confirmed in ccRCC models prior to therapeutic interpretation, ideally by direct measurement of cytosolic acetyl-CoA depletion and lipogenic flux. Protein SLC27A2 is present in our ccRCC proteomics datasets.

TYMP-high tumours clustered into networks associated with nucleotide metabolism, suggesting increased nucleoside turnover and salvage activity in ccRCC cells^42^. This is consistent with the broader metabolic reprogramming of ccRCC, in which tumour cells rely on altered metabolic pathways to sustain proliferation and survival^43,44^. In addition, TYMP has been reported to promote angiogenesis, inhibit apoptosis and modulate epigenetic regulation, and is overexpressed in several cancer types, including ccRCC^45^. Accordingly, tipiracil hydrochloride, a potent and selective TYMP inhibitor and component of trifluridine/tipiracil (TAS-102), which is approved for colorectal and gastric cancer, may represent a hypothesis-generating approach to limit ccRCC growth, although this requires dedicated functional validation^46^. Beyond these phenotype-defining candidates, VWF, LDHA, and TUBB supported exploratory secondary opportunities. VWF is a multimeric plasma glycoprotein that mediates platelet adhesion to the endothelium and participates in haemostasis and coagulation-related pathways^47^. Its elevated expression in several cancers suggests a role in tumour-associated vascular remodelling and prothrombotic signalling, processes relevant to ccRCC^47–51^, although no VWF-directed agent currently has an oncology indication and any therapeutic exploitation of this axis would require dedicated preclinical validation.

Among the tertiary-priority candidates, LDHA and TUBB emerged as the least compelling opportunities in our ccRCC cohort. LDHA is linked to glycolytic reprogramming, but the only FDA-approved LDHA-directed therapeutic, nedosiran, is a GalNAc-conjugated small interfering RNA that silences hepatic LDHA mRNA and is approved for primary hyperoxaluria type 1^52^; its biodistribution is deliberately restricted to hepatocytes, so direct clinical translation to renal tumour cells is not supported and it should be considered only as a conceptual prototype for LDHA silencing. TUBB-directed paclitaxel targeting also has limited rationale in ccRCC given the well-documented refractoriness of the disease to microtubule-targeting agents^53–56^.

In pRCC, we observed the same lipid metabolic reprogramming identified in ccRCC, supporting the view that this is a broader, pan-RCC metabolic feature^54–56^. ACLY emerged as a recurrent metabolic hub, being the protein with the highest overexpression in 60% of patients, consistent with reports showing that papillary renal tumours display increased mitochondrial activity and altered in situ metabolic characteristics and intracellular lipid features in pRCC^57,58^. Importantly, ACLY is the only top-priority target shared between ccRCC and pRCC in our data, positioning it as a candidate cross-subtype vulnerability and raising the possibility that an ACLY-directed strategy could be evaluated in a histology-agnostic, metabolism-defined subset of renal tumours, subject to the tissue-activation caveat described above.

TXNRD1-centred networks were enriched in cellular stress responses, NRF2-mediated nuclear events and ferroptosis^58,59^. This supported prioritisation of auranofin, an oral gold-containing drug indicated for rheumatoid arthritis^60^, as a hypothesis-generating candidate for TXNRD1-high pRCC. Mechanistically, auranofin has been reported to disrupt redox homeostasis and promote reactive oxygen species accumulation through TXNRD-related pathways^61^. In addition, auranofin combined with gemcitabine-based chemotherapy has been reported to enhance antitumour activity in pancreatic ductal adenocarcinoma^62^, suggesting that similar redox-directed strategies may merit evaluation in TXNRD1-high pRCC. In APOB-high pRCC cases, enrichment analysis highlighted cholesterol, fatty acid and lipoprotein metabolic pathways^55,56^, indicating a metabolic phenotype driven by altered lipid handling. Mipomersen, a liver-targeted second-generation antisense oligonucleotide previously approved for familial hypercholesterolaemia^63,64^, reduces circulating low-density lipoproteins by silencing APOB mRNA. Although its activity is restricted to the liver, mipomersen may serve as a conceptual prototype for lipid-directed metabolic intervention in APOB-high renal cancers, as systemic depletion of APOB-containing lipoproteins could limit the exogenous lipid and cholesterol supply available to these tumours. It is important to note, however, that mipomersen was withdrawn from the United States market in 2018 following persistent hepatotoxicity and injection-site reaction signals^65^, reinforcing that its role here is conceptual rather than a direct repurposing opportunity, and that translating APOB silencing into solid-tumour therapy will require liver-sparing delivery platforms.

PGD (6-phosphogluconate dehydrogenase) overexpression was also observed in selected pRCC cases. This key enzyme of the pentose phosphate pathway is frequently overexpressed across multiple cancer types and has been shown to promote tumour growth, metastatic potential and chemoresistance by supplying NADPH for anabolic processes and protection against oxidative stress^66–69^.

Integrating both RCC subtypes into a unified, top-three priority ranking (Fig. 4) provides a proof-of-concept for how proteomics-guided prescriptomics could be incorporated into future precision oncology workflows. By prioritising candidates according to TPA index and anchoring each nomination within its dominant pathway context, this framework moves beyond subtype-level suggestions towards patient-stratified therapeutic hypotheses. Crucially, these repurposing candidates are positioned as personalised coadjuvants to the prevailing standard of care, not as replacements for it, so that the clinician retains the current disease-appropriate backbone (ICI–TKI or MET-directed therapy for advanced ccRCC/pRCC as indicated) while gaining a molecularly justified, patient-specific add-on. These rankings are hypothesis-generating and require systematic functional validation, including target engagement and efficacy testing in appropriate preclinical models, before any clinical translation. Two of the therapeutic candidates, nedosiran and mipomersen, are GalNAc- or liver-restricted oligonucleotide therapeutics with negligible renal biodistribution, so their inclusion should be read as conceptual validation of the LDHA- and APOB-silencing axes rather than as direct repurposing opportunities; successful translation will require liver-sparing delivery systems. Bempedoic acid requires SLC27A2/ACSVL1-mediated activation, so the tumour-tissue expression of this activating enzyme should be confirmed before predicting ACLY engagement in renal tumours. Our approach should be interpreted in the context of the published proteogenomic RCC literature. The Clinical Proteomic Tumor Analysis Consortium (CPTAC) and related international efforts have profiled large cohorts spanning the principal and rare RCC subtypes by integrative proteogenomics. Clark *et al.* established the first comprehensive CPTAC clear cell RCC proteogenomic landscape, linking PBRM1/BAP1 status, immune infiltration patterns and metabolic rewiring to candidate therapeutic vulnerabilities^70^. Li *et al.* subsequently extended this framework to ccRCC aggressiveness, defining proteogenomic determinants of metastatic progression and outcome^71^. A complementary CPTAC effort then characterised rare RCC subtypes, including papillary and chromophobe RCC, providing the most directly comparable proteogenomic reference for the non-clear-cell tumours analysed here^72^. Qu *et al.* generated an independent proteogenomic ccRCC atlas in a Chinese patient cohort, complementing the CPTAC datasets with population-specific molecular features^73^. While these landmark studies typically emphasise relative quantification across hundreds of tumours and prioritise discovery of cohort-level drivers, our absolute, TPA-based approach is designed for within-patient stratification on a smaller, hypothesis-generating cohort. The two strategies are complementary: cross-cohort proteogenomic landscapes provide the disease-level reference frame and are powered to detect recurrent driver alterations, whereas absolute per-patient quantification provides the personalised dimension required for individual treatment selection. Tools that connect transcriptional or proteomic perturbation signatures to candidate compounds, such as the LINCS/Connectivity Map^74^, offer a further complementary layer that could be integrated with TPA-based prescriptomics in future work. Future studies should therefore explicitly benchmark TPA-based prescriptomics against these CPTAC-type proteogenomic resources, both to validate target prioritisation and to evaluate whether absolute quantification adds discriminative power beyond relative methods.

### Conclusions

This work establishes the TPA as a practical absolute-quantitation engine for personalised drug repurposing in renal cell carcinoma. By converting the per-patient proteome into a TPA index ranking and intersecting the resulting upregulated proteins with curated drug– target knowledge, the framework translates individual tumour biology, rather than histological subtype alone, into a structured shortlist of FDA-approved compounds positioned as molecularly justified coadjuvants to the prevailing standard of care. The same workflow accommodates a target-centric mode when an actionable protein crosses the prioritisation threshold and a pathway-centric mode when none does, so it remains usable across the heterogeneous biology of the principal RCC subtypes. Applied to 17 patient biopsies, the framework recovers biologically coherent and patient-specific drug nominations. In ccRCC, ACLY-high and TYMP-high patient subgroups support bempedoic acid and tipiracil hydrochloride, respectively, with VWF, LDHA, and TUBB as exploratory secondary options. In pRCC, the same TPA-driven prioritisation separates TXNRD1-high, ACLY-high and APOB-high patients, mapping onto auranofin, bempedoic acid and mipomersen as primary candidates. Each nomination is anchored to a measurable, patient-specific abundance changes rather than to a population average. These results constitute a proof of concept rather than a clinical recommendation.

## Acknowledgements

This work was funded by national funds from FCT - Fundação para a Ciência e a Tecnologia, I.P., under the scope of the projects UID/50006/2025, UID/PRR/50006/2025, UID/PRR2/50006/2025 and LA/P/0008/2020 of the Associated Laboratory for Green Chemistry - LAQV REQUIMTE (https://doi.org/10.54499/UID/50006/2025, https://doi.org/10.54499/UID/PRR/50006/2025, https://doi.org/10.54499/UID/PRR2/50006/2025, and https://doi.org/10.54499/LA/P/0008/2020). H.M.Santos acknowledges his research contract that is funded by national funds from FCT – *Fundação para a Ciência e a Tecnologia*, I.P., under the scope of the project LA/P/0008/2020 of the Associated Laboratory for Green Chemistry - LAQV REQUIMTE (DOI: 10.54499/LA/P/0008/2020). A.Q.F. is funded by the FCT, I.P, PhD grant (2023.00528.BD). I.F.D. is funded by the FCT/MCTES PhD grant (2024.00745.BD). PROTEOMASS Scientific Society is acknowledged by the funding provided to the Laboratory for Biological Mass Spectrometry – *Isabel Moura* (#PM001/2024, #PM001/2019 and #PM003/2016).

## Authors’ Contributions

R.D. collected and organised all human tissue biopsies. J.L.C., C.L., H.M.S., G. M., D. K. and R.D. designed the re-analysis framework and provided financial support. A.Q.F. performed the proteomics data processing and TPA-based quantification under the supervision of H.M.S and J.L.C. A.Q.F., and I.F.D. performed the bioinformatic analyses under the supervision of H.M.S and J.L.C. D.K. and J.L.C. drafted the first manuscript. C.L., R.D., L.B.C., L.M., J.R.W., G.M., M.M., L.C.P., H.M., D.K., H.M.S. and J.L.C. revised the draft and provided critical input or scientific comments. All authors read and approved the final version of the manuscript.

## Funding

This work was funded by national funds from FCT - Fundação para a Ciência e a Tecnologia, I.P., under the scope of the projects UID/50006/2025, UID/PRR/50006/2025, UID/PRR2/50006/2025 and LA/P/0008/2020 of the Associated Laboratory for Green Chemistry - LAQV REQUIMTE (https://doi.org/10.54499/UID/50006/2025, https://doi.org/10.54499/UID/PRR/50006/2025, https://doi.org/10.54499/UID/PRR2/50006/2025 and https://doi.org/10.54499/LA/P/0008/2020). PROTEOMASS Scientific Society provided funding through the general funding programmes #PM001/2024, #PM001/2019, #PM003/2016. A.Q.F. is supported by FCT PhD grant 2023.00528.BD; I.F.D. is supported by FCT/MCTES PhD grant 2024.00745.BD; H.M.S.’s research contract is funded under LA/P/0008/2020. The University of Pittsburgh Hillman Cancer Center shared resource facilities were supported in part by NIH/NCI award P30CA047904.

## Declarations

### Ethics approval and consent to participate

The study was approved by the Institutional Review Board at the University of Pittsburgh (IRB #02-077).

### Consent for publication

Not applicable.

### Competing interests

The authors declare no potential conflicts of interest.

## Author details

### Availability of data and materials

The mass spectrometry proteomics data have been previously deposited to the ProteomeXchange Consortium via the PRIDE^36^ partner repository with the dataset identifier PXD023296.

## Abbreviations

ACLY: ATP-Citrate Synthase
AKT: Protein Kinase B
APOB: Apolipoprotein B-100
BAP1: BRCA1-Associated Protein-1
CAIX: Carbonic Anhydrase 9
ccRCC: Clear Cell Renal Cell Carcinoma
COMT: Catechol O-methyltransferase
CPTAC: Clinical Proteomic Tumor Analysis Consortium
CTLA-4: Cytotoxic T-Lymphocyte-Associated Protein 4
DNA: Deoxyribonucleic Acid
FDA: Food and Drug Administration
FH: Fumarase Hydratase
HIF-1α: Hypoxia-Inducible Factor 1α
HIF-2α: Hypoxia-Inducible Factor 2α
HLRCC: Hereditary Leiomyomatosis and Renal Cell Carcinoma
ICI–TKI: Immune Checkpoint Inhibitors - Tyrosine Kinase Inhibitors
IMPDH2: Inosine-5’-monophosphate dehydrogenase 2
IRB: Institutional Review Board
KEGG: Kyoto Encyclopedia of Genes and Genomes
LC-MS/MS: Liquid Chromatography–Tandem Mass Spectrometry
LDHA: L-lactate Dehydrogenase A Chain
MET: Mesenchymal-Epithelial Transition
mTOR: Mechanistic Target of Rapamycin
NADPH: Nicotinamide Adenine Dinucleotide Phosphate
NAT: Normal Adjacent Tissue
NFE2L2: Nuclear Factor, Erythroid 2 Like 2
OCT: Optimal Cutting Temperature
PARP1: Poly [ADP-ribose] Polymerase 1
PBRM1: Polybromo-1
PD-1: Programmed Cell Death Protein 1
PDGF: Platelet-derived Growth Factor
PGD: 6-Phosphogluconate Dehydrogenase
PI3K: Phosphoinositide 3-Kinase
PLIN2: Perilipin-2
pRCC: Papillary Renal Cell Carcinoma
RCC: Renal Cell Carcinoma
RNA: Ribonucleic Acid
SLC27A2/ACSVL1: Solute Carrier Family 27 Member 2
TAS-102: Chemotherapy Drug Trifluridine/ Tipiracil
TCA: Citric Acid Cycle
TKI: Tyrosine Kinase Inhibitors
TPA: Total Protein Approach
TTD: Therapeutic Target Database
TUBB: Tubulin Beta Chain
TUBB3: Tubulin beta-3 chain
TXNRD1: Thioredoxin Reductase 1, Cytoplasmic
TYMP: Thymidine Phosphorylase
VEGF: Vascular Endothelial Growth Factor
VEGFR: Vascular Endothelial Growth Factor Receptor
VHL: Von Hippel–Lindau
VWF: Von Wilebrand Factor

## Notes

### Competing Interest Statement

The authors have declared no competing interest.

