## Supplementary figures and images for "Absolute quantitative proteomics guides patient-stratified drug repurposing in clear cell and papillary renal cell carcinoma"

### Supplementary Figure 1.1

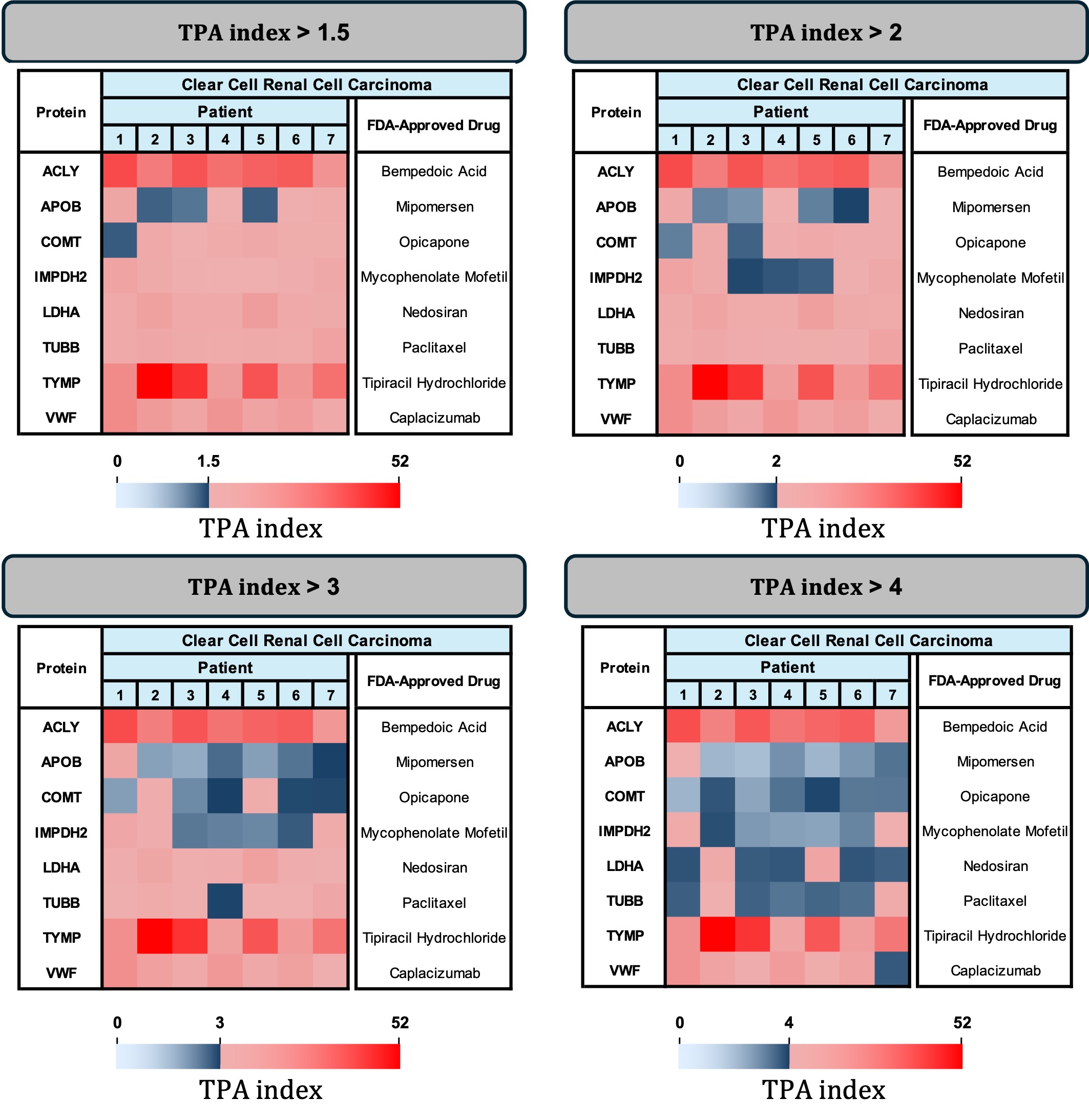

### Supplementary Figure 1.2

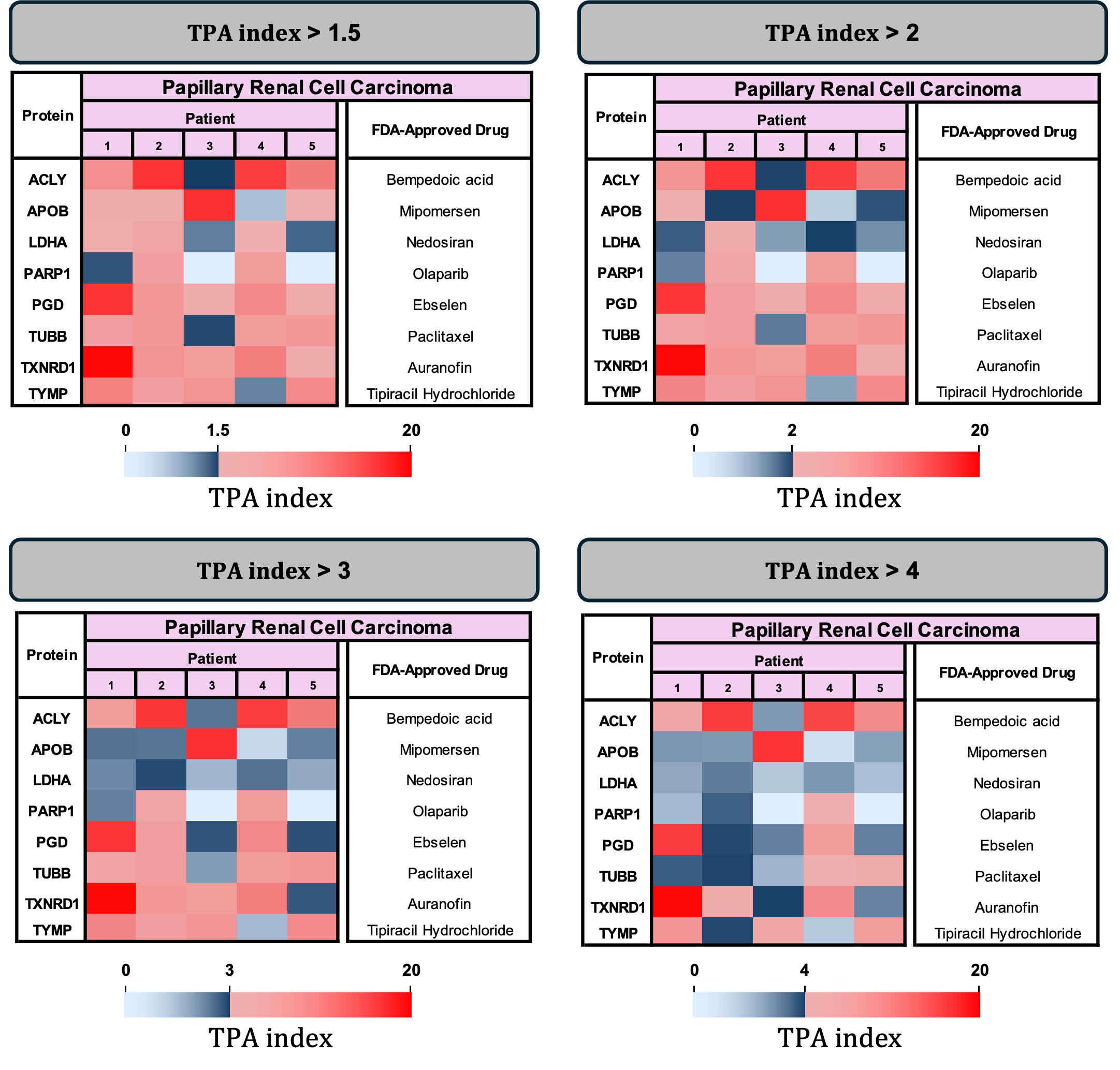
