## Supplementary Material - Figures for "Absolute quantitative proteomics guides patient-stratified drug repurposing in clear cell and papillary renal cell carcinoma"

**Supplementary Table 1**: Clinical and pathological characteristics of human kidney biopsies included in this study.

| **Biospy** | **ID** | **Gender** | **Age** | **Sample Type** | **Diagnosis** |
| --- | --- | --- | --- | --- | --- |
| **N1** | N1 | M | 54 | NAT | RCC |
| **N2** | N2 | F | 49 | NAT | pRCC |
| **N3** | N3 | F | 58 | NAT | RCC |
| **N4** | N4 | F | 72 | NAT | RCC |
| **N5** | N5 | M | 70 | NAT | RCC |
| **CC21** | **1** | M | 50 | ccRCC | RCC |
| **CC22** | **2** | M | 68 | ccRCC | RCC |
| **CC23** | **3** | M | 60 | ccRCC | RCC |
| **CC24** | **4** | F | 68 | ccRCC | RCC |
| **CC25** | **5** | M | 58 | ccRCC | RCC |
| **CC26** | **6** | F | 42 | ccRCC | RCC |
| **CC27** | **7** | F | 58 | ccRCC | RCC |
| **P16** | **1** | F | 51 | pRCC | RCC |
| **P17** | **2** | M | 66 | pRCC | RCC |
| **P18** | **3** | M | 75 | pRCC | RCC |
| **P19** | **4** | M | 87 | pRCC | RCC |
| **P20** | **5** | M | 69 | pRCC | RCC |

**M**: Male; **F**: Female; **RCC**: Renal cell carcinoma **NAT**: Normal adjacent tissue; **ccRCC**: Clear cell renal cell carcinoma; **chRCC**: Chromophobe renal cell carcinoma; **pRCC**: Papillary renal cell carcinoma

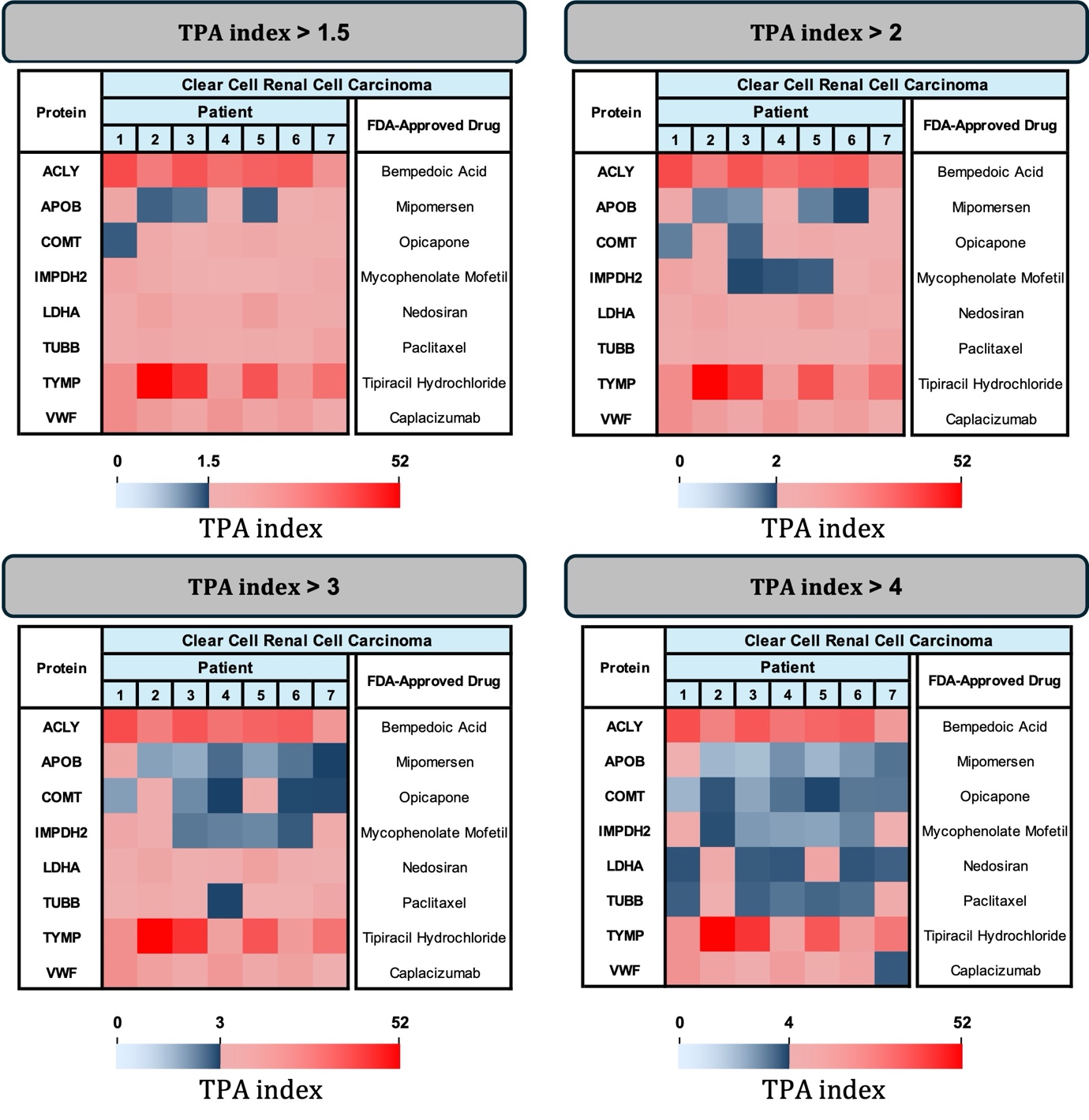
**Supplementary Figure 1**: Comparison of therapeutic target prioritisation in ccRCC patients across different selection thresholds. FDA-approved drugs targeting eight upregulated proteins are shown for seven patients at fold-change thresholds of 1.5, 2 (the threshold applied in the main text), 3, and 4.

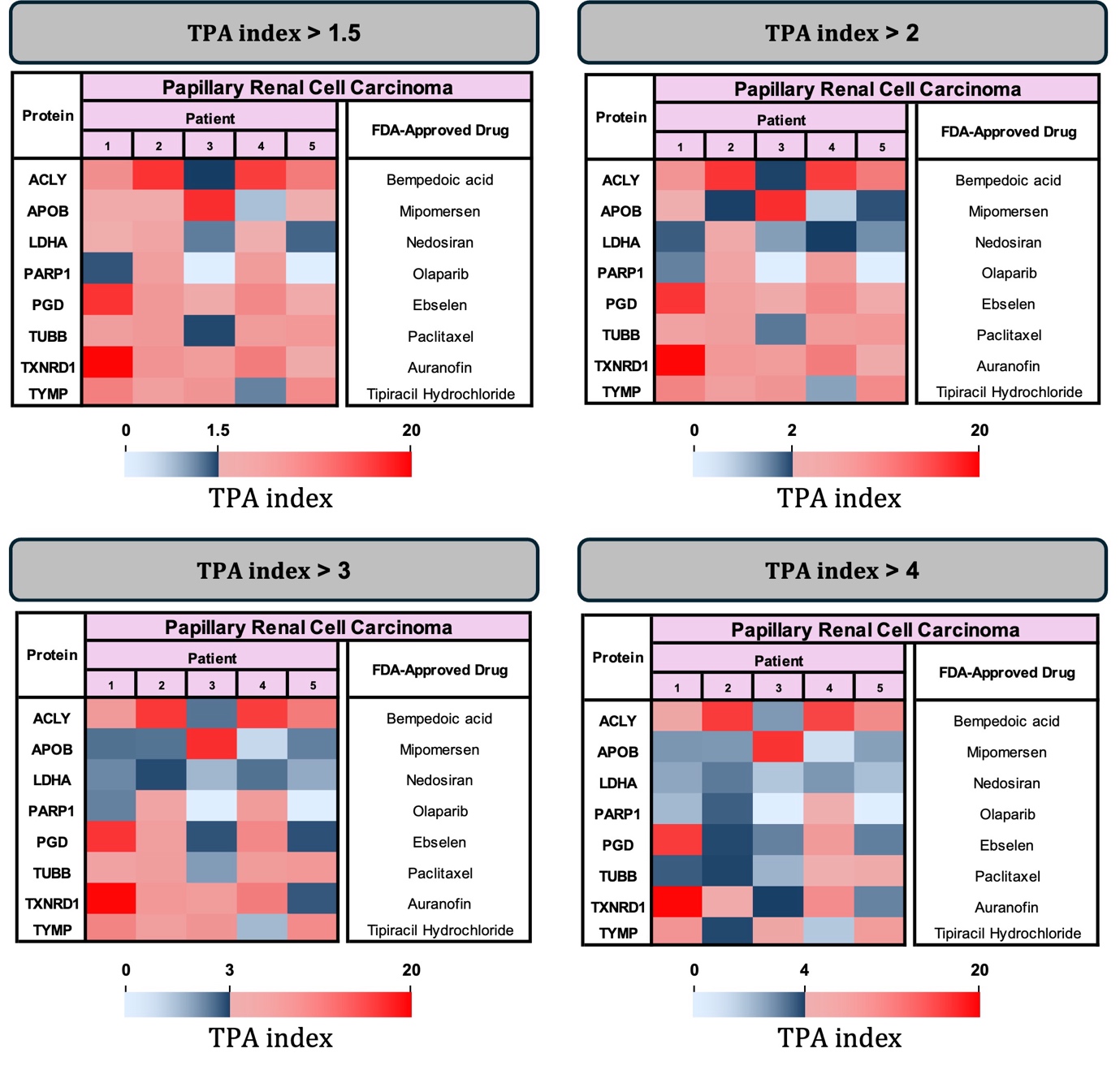

**Supplementary Figure 2**: Comparison of therapeutic target prioritisation in pRCC patients across different selection thresholds. FDA-approved drugs targeting eight upregulated proteins are shown for seven patients at fold-change thresholds of 1.5, 2 (the threshold applied in the main text), 3, and 4.

**Supplementary Discussion**

In both RCC subtypes, several candidate targets maintain high $TPA index$ values even under In both RCC subtypes, most candidate targets retain high $TPA index$ values even as increasingly stringent log_2_ fold-change thresholds are applied, indicating stable and marked overexpression at the individual‑patient level. Across cut-offs thresholds, the top three proteins selected per patient (Fig. 4) remain largely unchanged, only papillary RCC patients 2 and 3 lose a single indication each (TUBB and TXNRD1, respectively), suggesting that targets which disappear at higher cut‑offs represent context‑dependent or weaker signals rather than robust candidates. Quantitatively, the proportion of retained indications in pRCC remains substantial, 80%, 67.5%, 55% and 40% at log_2_ fold‑change thresholds of 0.5, 1, 1.5 and 2, respectively. For ccRCC a higher stability is shown with 93%, 84%, 71% and 48% retention at the same thresholds. Together, these observations support the utility of the TPA‑based framework for differentiating durable, patient‑level therapeutic opportunities from less reliable hits that are sensitive to arbitrary fold‑change choices.
